# DNA Methylome Regulates Virulence and Metabolism in *Pseudomonas syringae*

**DOI:** 10.1101/2024.02.12.579912

**Authors:** Jiadai Huang, Fang Chen, Beifang Lu, Yue Sun, Youyue Li, Canfeng Hua, Xin Deng

## Abstract

Bacterial pathogens employ epigenetic mechanisms, including DNA methylation, to adapt to environmental changes, and these mechanisms play important roles in various biological processes. *Pseudomonas syringae* is a model phytopathogenic bacterium, but its methylome is less well known than that of other species. In this study, we conducted single-molecule real-time sequencing to profile the DNA methylation landscape in three model pathovars of *P. syringae*. We identified one Type-I restriction-modification system (HsdMSR), including the conserved sequence motif associated with N^6^-methyladenine (6mA). About 25%–40% of the genes involved in DNA methylation were conserved in two or more of the strains, revealing the functional conservation of methylation in *P. syringae*. Subsequent transcriptomic analysis highlighted the involvement of HsdMSR in virulent and metabolic pathways, including the Type III secretion system, biofilm formation, and translational efficiency. The regulatory effect of HsdMSR on transcription was dependent on both strands being fully 6mA methylated. Overall, this work illustrated the methylation profile in *P. syringae* and the critical involvement of DNA methylation in regulating virulence and metabolism. Thus, this work contributes to a deeper understanding of epigenetic transcriptional control in *P. syringae* and related bacteria.

## Introduction

*Pseudomonas syringae* inflicts leaf spots and cankers on plants globally, and its adequate control poses significant challenges. More than 60 pathovars have been identified, which infect almost all economically important crops in the world (1, 2). *P. syringae* pv. *phaseolicola* 1448A (*Psph*), one of the classical strains of *P. syringae*, can cause severe halo blight of common beans (*Phaseolus vulgaris*), which raises it from a common pathogen to a molecular plant-pathogen bacterium(3). *P. syringae* pv. *tomato* DC3000 (*Pst*) and *P. syringae* pv. *syringae* B728a (*Pss*) are two other model strains whose natural host plants are tomatoes and beans, respectively (4, 5). The most important weapon of *P. syringae,* and the first to be characterised, is the Type III secretion system (T3SS) encoded by *hrp* and *hrc* gene clusters, which are flanked by the conserved effector locus and deliver T3 effectors into host cells (6).

The expression of T3SS genes is inhibited in nutrient media such as King’s B (KB), whereas it is induced in minimal medium or plant cells (7–9). The HrpRSL pathway is the primary regulator of the T3SS in *P. syringae*, with HrpRS activating *hrpL* expression. HrpL then binds to the *hrp* box to induce the translation of downstream T3 effectors (10–12). HrpRSL also impacts other virulence-related mechanisms, including motility, biofilm formation, siderophore production, and oxidative stress resistance, which are under the control of complicated regulatory networks, allowing *P. syringae* to infect plants (13–19). However, the methylome and function of DNA methylation in *P. syringae* pathogenesis and metabolism remain largely unknown.

In bacteria, DNA methylation is the primary level of epigenetic regulation because these prokaryotes lack the histones and nucleosomes of eukaryotic cells. Bacterial DNA has three primary forms of methylation: N^6^-methyladenine (6mA), N^4^-methylcytosine (4mC), and N^5^-methylcytosine (5mC). The latter is the most common type in eukaryotes, while 6mA is the dominant form in prokaryotes. DNA methylation in bacteria is mainly the product of methyltransferase (MTase) enzymes, which transfer a methyl group from S-adenosyl-l-methionine to various positions on target bases, depending on the modification (20). MTases originate from the restriction-modification (R-M) system, which protects bacterial cells from invading DNA by recognising and cleaving specific unmethylated motifs (21, 22). There are three main types of R-M systems, Types I, II, and III, categorised according to the related subunits and the precise site of DNA restriction (23, 24). Additionally, orphan MTases are an emerging group of MTases without cognate restriction that are involved in the regulation of DNA replication and gene expression (21). The Type I R-M system contains three host-specificity determinant (*hsd*) subunits, restriction (R), modification (M), and specificity (S), which are encoded by *hsdR*, *hsdM*, and *hsdS*, respectively. Genome sequencing and bioinformatics analyses have revealed that the HsdMSR system exists in around half of all bacteria and archaea species (24). Nonetheless, the biological roles and specific motifs alongside their targets of most MTases remain unknown, especially those related to 6mA (25).

A new sequencing technology, single-molecule real-time (SMRT) sequencing, can detect all three forms of DNA methylation, but particularly 6mA (26). SMRT-seq has been applied for complete genome sequencing and insertion sequence profiling in different *P. syringae* pv. *actinidiae* (*Psa*) strains (27–29). In *P. aeruginosa*, Type I R-M systems, along with their specific motifs, have been identified in the model strain PAO1 and two clinical strains via SMRT-seq. These MTases are considered to play important roles in *P. aeruginosa* virulence or drug resistance (30–32).

To study the targets and functions of DNA modifications in *P. syringae*, SMRT-seq was used to identify global methylation sites and reveal their conserved and divergent functions in the three model strains in this study. We found that HsdMSR in *Psph* was involved in modulating three pathways, namely T3SS, biofilm production, and the metabolism-related gene expression of ribosomal proteins. Moreover, we found that HsdMSR-mediated transcriptional regulation depends on the full methylation of both DNA strands. Taken together, our results provide insights into the involvement of DNA methylation in the phenotypic traits of *P. syringae* and transcription regulatory mechanisms that may apply to other pathogenic bacteria.

## Results

### Genome-wide methylome profiling of three model *P. syringae* strains

According to REBASE (33) database prediction, 0 to 2 Type I R-M systems and 3 to 4 Type II R-M systems exist in the three pathovars of *P. syringae*, indicating that Type I is more conserved than Type II in this pathogen **(Table S1)**. To characterise their methylomes, we performed SMRT-seq to obtain the methylation patterns on a genome-wide scale using DNA extracted from the stationary phase of the three wild-type (WT) strains. Our results revealed a total of 10,302 modified bases, of which 3,849, 4,646, and 1,807 nucleotides were significantly identified as being 6mA, 5mC, and 4mC modified, with an interpulse duration ratio (IPD) of > 1.5 throughout the genome of *Psph* **(Fig. 1A)**. After identifying the significantly modified sites, we orientated the gene locus tags within the methylated bases. Most genes harboured only 6mA modifications, although 339 genes harboured all three DNA methylation types **(Fig. 1B)**.

**Figure 1.**
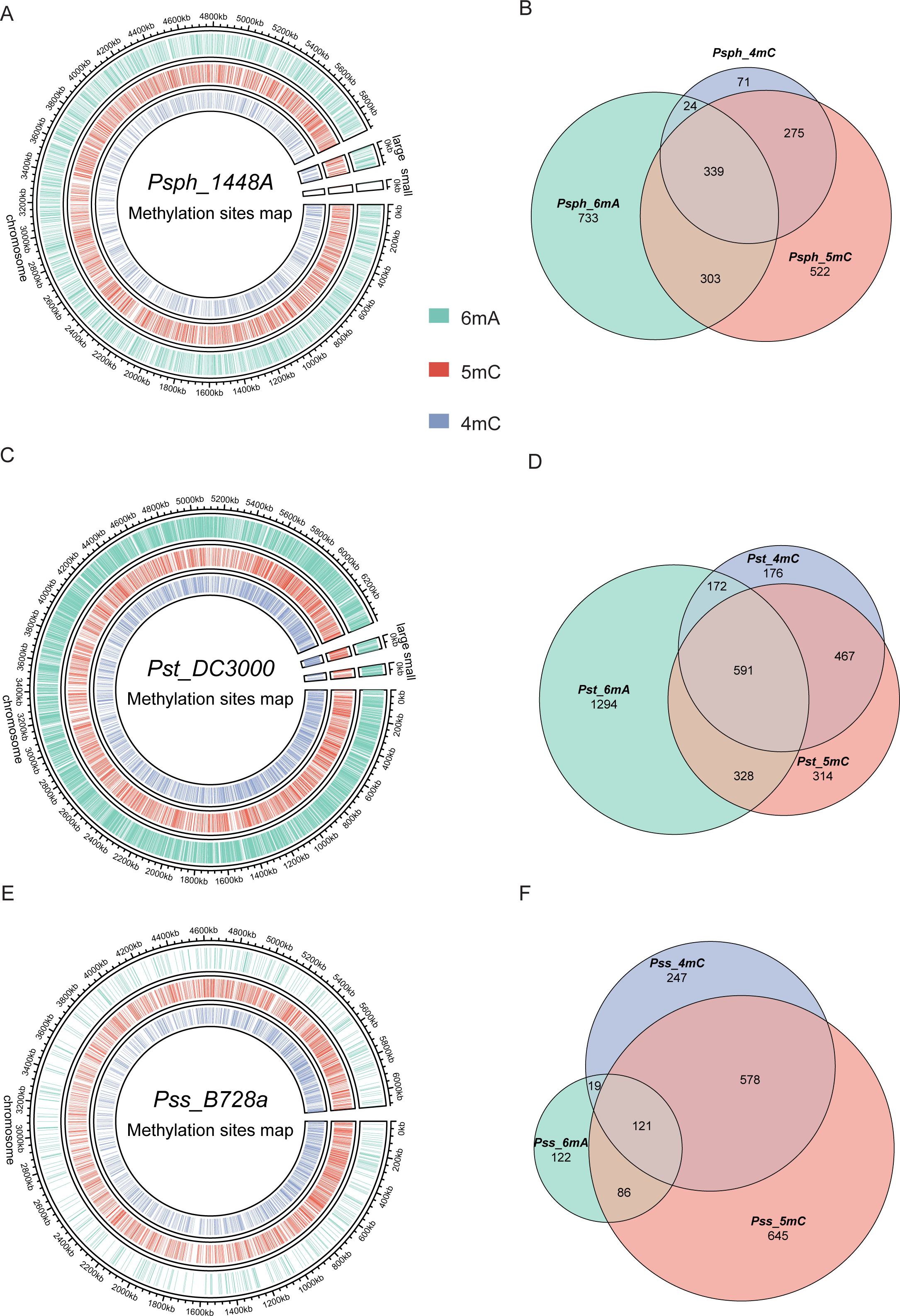
Genome-wide identification of *P. syringae* DNA methylation. **(A)** The circle map displays the distribution of 6mA, 5mC, and 5mC in *Psph* WT. **(B)** The Venn plot reveals overlapped genes within three types of DNA methylation of *Psph* WT. **(C)** The circle map displays the distribution of 6mA, 5mC, and 5mC in *Pst* WT. (**D**) The Venn plot reveals overlapped genes within three types of DNA methylation of *Pst* WT. **(E)** The circle map displays the distribution of 6mA, 5mC, and 5mC in *Pss* WT. **(F)** The Venn plot reveals overlapped genes within three types of DNA methylation of *Pss* WT.

In *Pst*, a total of 6,002, 2,242, and 3,514 methylation sites were significantly identified as 6mA, 4mC, and 5mC, respectively, using the same cut-off (IPD >1.5) **(Fig. 1C)**. Among these modified bases, more genes equipped with all three modification types were detected in *Pst* than *Psph*, with 591 (17.7%) genes exhibiting modifications in *Pst* sequences **(Fig. 1D)**. Additionally, the methylome atlas of *Pss* revealed a lower incidence of methylation than those of *Psph* and *Pst*, particularly in terms of 6mA modifications, which were only seen in 457 significant 6mA occurrences under the same threshold (IPD > 1.5) and a total of 2,853 and 1,438 methylation sites were detected as 5mC and 4mC, respectively **(Fig. 1E)**. Comparative analysis of the three methylations in *Pss* showed that the highest numbers of genes were modified by 4mC and 5mC simultaneously (**Fig. 1F)**.

### Characteristics of modified loci and GC contents in *P. syringae*

To further uncover the distribution of the modifications throughout the genome, we calculated how many of the three kinds of modified sites were in these three strains. The majority of modifications occurred in coding sequence (CDS) regions, accounting for at least 80% **(Fig. S1A)**, and less than 20% were in intergenic regions and non-coding RNA (tRNA and rRNA). However, compared with 4mC and 5mC, 6mA was more likely to be in intergenic regions, which suggests that it potentially functions in transcription regulation in *P. syringae*.

It is known that 5mC occurs within CpG sites in CpG islands, which contain higher GC percentages than is typical in the human genome (34). Additionally, the GC architecture can influence DNA methylation in eukaryotic cells (35). To determine the GC content characteristics of modification sites in *P. syringae*, we extracted the sequences 50-bp upstream and downstream from the modified bases and calculated their GC content. The distribution of modification sites and their GC content are shown in density plots. Compared with 4mC and 5mC, the GC contents of 6mA sites were the lowest and the closest to the average GC percentage of *P. syringae* (58% for *Psph* and *Pst*, 59.2% for *Pss*) **(Fig. S1B)**. In contrast, 5mC modification bases had the highest GC content, especially in *Pss* **(Fig. S1C)**. Furthermore, 4mC sites in *Psph* had a higher GC content than those in *Psa* and *Pss* **(Fig. S1D)**. Similar phenomena have been observed in various other bacterial species. For instance, the *Escherichia coli* MTase Dcm can catalyse the 5′CCWGG3′ motif (36). Furthermore, in *Spiroplasma* sp. strain MQ-1, more than 95% of 5mC modifications are found in high-CG sequences, which is similar to the pattern in eukaryotes (37).

### Methylated genes are functionally conserved among three *P. syringae* strains

When we compared the three methylation types in the three tested strains, we observed the conservation pattern among them **(Fig. 2A-B and Table S2)**. Notably, about 25% to 45% of methylated genes were conserved in two and three strains. Interestingly, 5mC had the highest conservation level (39.1%), which might be explained by the rare occurrence of 6mA in *Pss*. Conversely, an obviously higher degree of conservation was observed in virulence-related genes, including T3SS and alginate biosynthesis-related genes, between *Psph* and *Pst* **(Table S2)**. Additionally, some methylated genes (n = 739) harboured sites for different modification types in the three strains **(Fig. 2A)**. For example, the modification sites of a Cro/CI family transcriptional factor (TF) in PSPPH_1319 were for different modification types in the three *P. syringae* strains. The Cro/CI are important TFs present in phages. The interaction between Cro and CI affects bacteria immunity status in Enterohemorrhagic *Escherichia coli* (EHEC) strains (38). *Psph* carried 6mA and 5mC, while *Pst* and *Pss* had 5mC and 4mC. This difference might result from differences in the MTases and their specific motifs **(Table S2)**.

**Figure 2.**
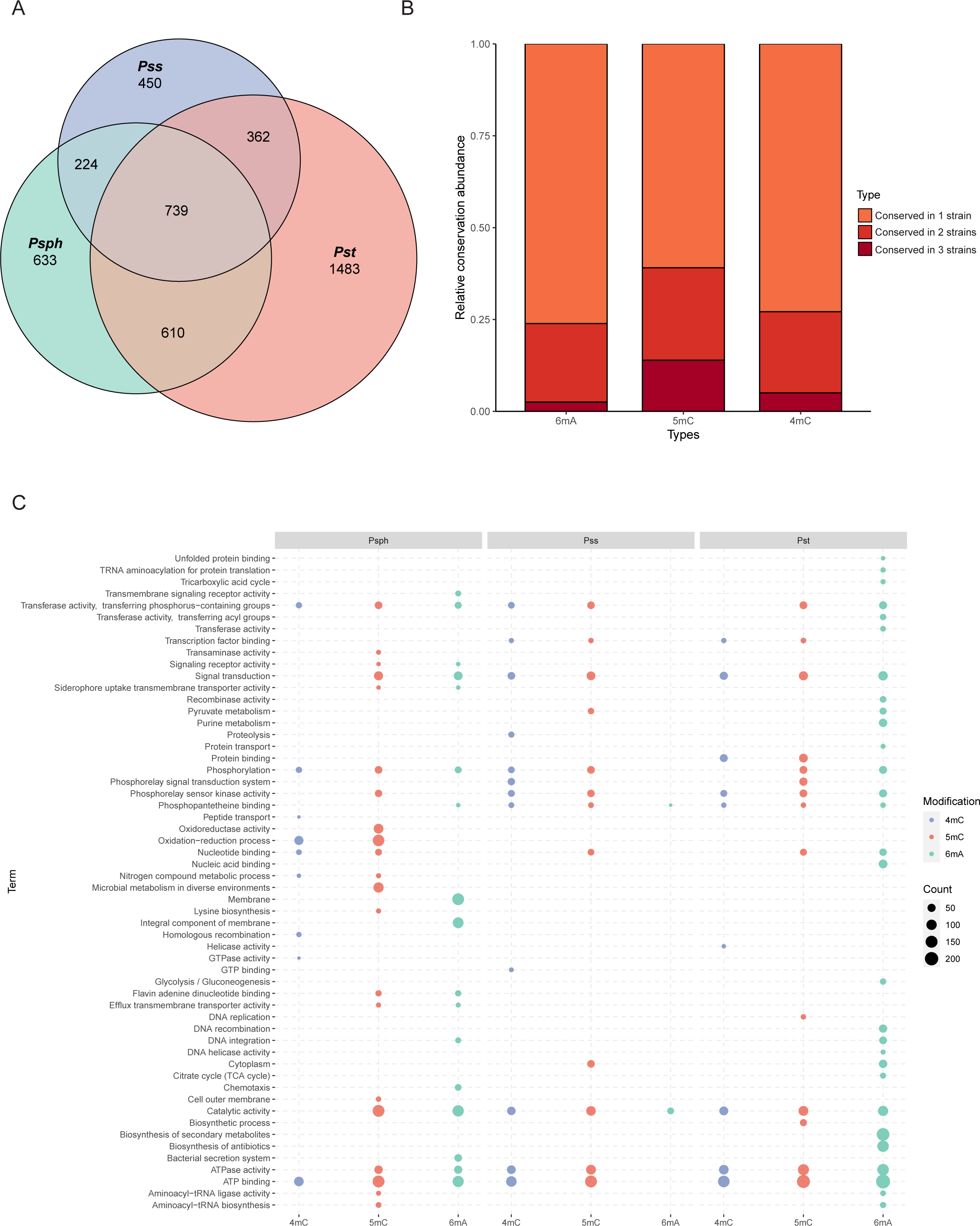
Functional enrichment analysis of methylation sites in three *P. syringae* strains. **(A)** Repartition of the total pool of modified genes among strains. **(B)** Proportion of methylated genes detected in one, two, or three genomes for all *P. syringae* strains and conserved DNA methylation sites with detected genes. **(C)** The dot plot revealed the significantly enriched functional pathways in GO and KEGG databases among three *P. syringae* strains. The specific names of each pathway were listed on the left, and each column with dots indicated the number of genes within one kind of methylation in one of three *P. syringae* strains. The size of the dots indicates the number of related genes.

To elucidate the functions of genes methylated with these three modification types, functional enrichment analyses were performed based on gene ontology (GO) and Kyoto Encyclopedia of Genes and Genomes (KEGG) databases (39, 40). Analysis revealed the shared functional characteristics of genes with 5mC and 4mC modifications in *Pss* and *Pst*. In contrast, genes with 6mA modifications exhibited more conserved functional terms between *Psph* and *Pst* **(Fig. 2C)**. For instance, *Pst* and *Pss* contained 4mC-methylated genes enriched in signalling transduction and TF binding, whereas *Psph* exhibited enrichment in oxidation–reduction process and nucleic acid binding genes. Remarkably, notable conservation of functional terms was observed for genes with 6mA modifications, with at least 21 terms conserved between *Psph* and *Pst*. These terms included ATP binding, catalytic activity, integral component of membrane, transferase activity, and phosphorylation. We also conducted Cluster of Orthologous Group (COG) protein function analysis (41, 42), which assigned the methylated genes from the three strains to 20 categories with diverse functions **(Fig. S2A-C)**. In *Psph*, the top three COG classifications encompassed inorganic ion transport, amino acid transport, and transcription **(Fig. S2A)**. Furthermore, the abundance of modified genes related to cell wall/membrane/envelope biogenesis (M category) in *Pss* was lower than that of *Psph* and *Pst*. Despite the slight variations in COG distribution, the overall pattern exhibited considerable similarity and conservation across the three strains.

### Five conserved sequence motifs of 6mA and 4mC were identified in *P. syringae*

Obtaining the whole methylomes of the three strains led us to investigate the associated MTases and their target motifs. The PacBio motif finder and MEME suite (43) were used to determine the specific sequence motifs within 50 bp upstream and downstream of the significantly modified bases. Using the results of both software packages with corresponding cut-offs, we identified two 6mA motifs in *Psph:* C**A**GCN(_6_)CTC and RAGT**A**CTY **(Fig. 3A)**. The bolded bases indicate the methylation sites in these motifs. The two motifs occurred a total of 2,998 and 300 times in double strands, and more than 98% and 77% of those occurrences were methylated, respectively, revealing the high methylation rates of these two motifs and implying their crucial roles in *Psph* **(Fig. 3B)**. However, the analysis did not obtain any identified motifs for cytosine modifications.

**Figure 3.**
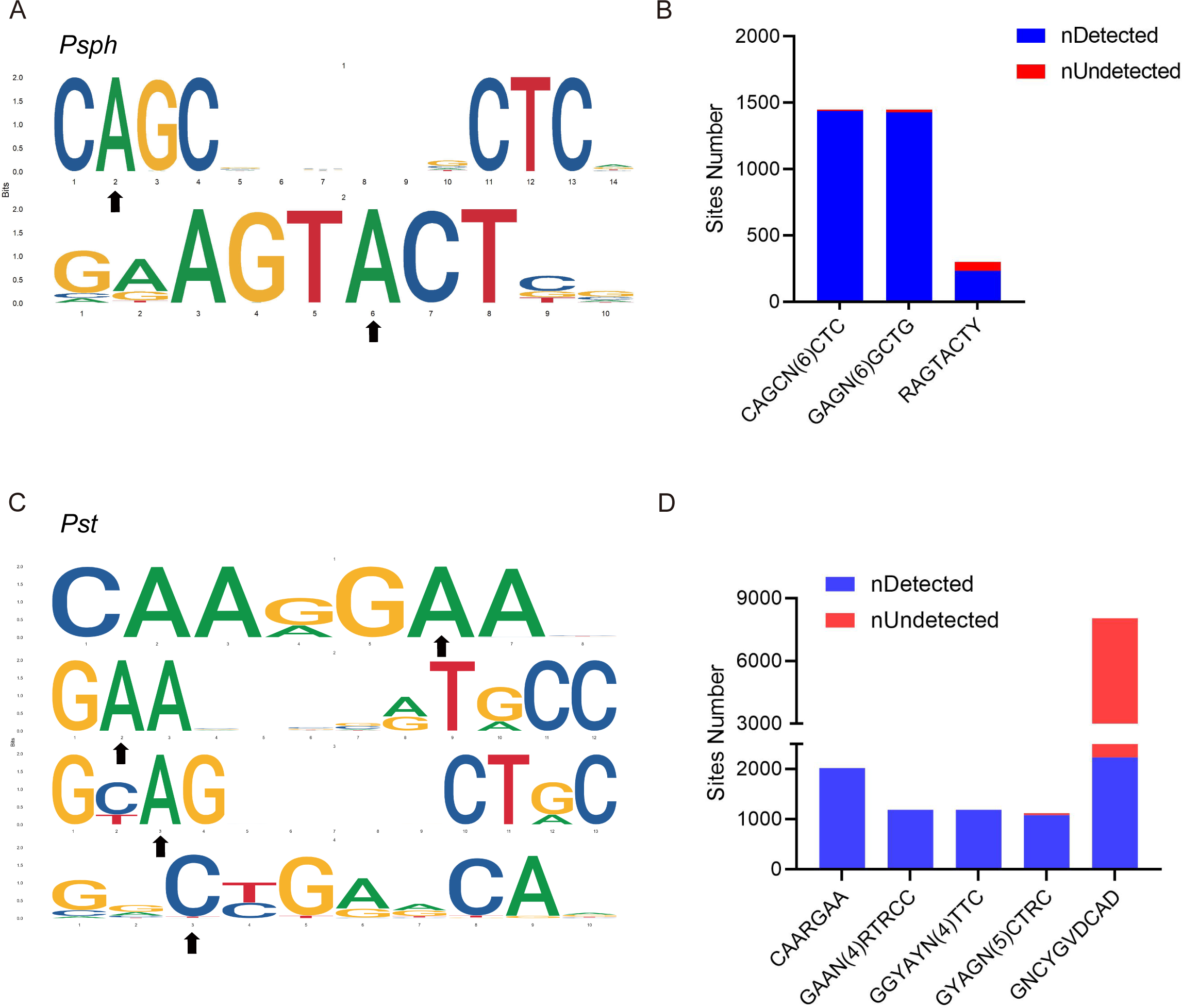
DNA methylation motifs in *P. syringae*. **(A)** 6mA methylation motifs found in *Psph* using SMRT-seq. The black arrows indicate the site of adenine methylation. **(B)** Bar plot shows the abundance of methylated numbers throughout all motif sites in *Psph*. **(C)** 6mA and 4mC methylation motifs found in *Pst* using SMRT-seq. The black arrows indicate the site of base methylation. **(D)** Bar plot shows the abundance of methylated numbers throughout all motif sites in *Pst*.

Three motifs for 6mA and one motif for 4mC were identified in *Pst* **(Fig. 3C)**. The first 6mA motif, CAARG**A**A, was found 2,022 times in the genome, with 2,019 of the occurrences methylated (99.8%) **(Fig. 3D)**. The second 6mA motif, G**A**AN(_4_)RTRCC, was fully methylated in both strands. Similarly, the last 6mA motif, GY**A**GN(_5_)CTRC, exhibited a high methylation level (96%) in the genome of *Pst*. Conversely, the only motif identified for 4mC in *Pst* displayed a considerably lower methylation level than the 6mA sites, with 2,207 occurrences of methylation out of a total of 8,048 sites (approximately 27%). Notably, no credible motifs were found in *Pss* because of its noticeably lower modification levels compared with the other two strains. Overall, the application of SMRT-seq to the three model *P. syringae* strains identified specific sequence motifs, including 6mA motifs, exhibiting extensive methylation statuses.

### *In vivo* validation revealed the activity and specificity of Type I R-M system MTase in *Psph*

The first motif, C**A**GCN(_6_)CTC, identified in *Psph* is similar to the counterpart modified by a Type I R-M MTase in *P. aeruginosa* (26) **(Fig. S3)**, suggesting that the putative HsdMSR is responsible for the observed motif. As no other motifs exhibited a discernible association with the Type II R-M MTases, we opted to focus our investigation on the putative HsdMSR in *Psph.* To determine the potential functions of the putative MTases, we constructed MTase knock-out mutants and complementary strains of HsdMSR. Growth curve experiments showed the adverse effects on Δ*hsdMSR* compared to the WT strain **(Fig. S4A)**. To confirm the activity of HsdMSR, a dot blot assay was used to detect the 6mA intensity in WT and Δ*hsdMSR*. The results shown in **Figure S4B** illustrated that 6mA levels were significantly decreased in Δ*hsdMSR* compared to the WT, but were restored in the complemented strain. In addition, SMRT-seq was performed on Δ*hsdMSR*, which further profiled methylation patterns in Δ*hsdMSR* **(Fig. S4C-D)**. As expected, we found almost all of the reduction 6mA sites in the Δ*hsdMSR* were from motif CAGCN(6)CTC **(Fig. S4E-F)**. We also noticed that 5mC and 4mC sites in the mutant were relatively similar to that in WT **(Fig. S4E),** and the slight difference might be caused by sequencing errors. Overall, we propose that HsdMSR only catalyze the 6mA located on the motif CAGCN(6)CTC, but does not affect other 6mA sites and other modification types.

In accordance with previous studies showing that growth conditions affect the bacterial methylation status (44–46), we applied dot blot experiments using the same amount of DNA (1 μg) from these three *P. syringae* strains to detect the 6mA levels during both logarithmic and stationary phases. The results revealed that 6mA levels in the stationary phase were much higher than those in the logarithmic phase in *Psph* and *Pst*, but no significant change in *Pss* **(Fig. S5A)**. Additionally, we found that during the stationary phase, 6mA methylation levels in *Psph* and *Pst* were higher than those in *Pss*. These findings were consistent with the MTases predication on these three strains since *Pss* does not harbor any type I R-M systems, which are important for 6mA medication in bacteria. We also overexpressed HsdM in *Pst* and performed additional experiments in WT and the HsdM overexpression strains, including dot blot and growth curve assays. The results showed that the MTase overexpressed strain presents a higher 6mA level than WT during the logarithmic phase, and the overexpression of MTase had no effects on growth in *Pst* **(Fig. S5B-C)**. Taken together, the results demonstrated that 6mA levels change with the bacterial growth phase in *Psph* and *Pst*, and HsdMSR is responsible for maintaining 6mA sites within the sequence motif of C**A**GCN(_6_)CTC in *Psph*.

### Transcriptomic analysis profiling of differentially regulated genes associated with virulence and metabolism in the HsdMSR mutant

To explore the regulatory influence of HsdMSR on gene expression, RNA-seq was applied to analyse WT and Δ*hsdMSR* in the stationary growth phase, which ensured that the strains had sufficient DNA methylation levels. Differentially expressed genes (DEGs) were obtained with the cut-off of |log_2_FC| > 1 and an adjusted p-value < 0.05. We identified 395 DEGs between WT and Δ*hsdMSR* under these experimental conditions. Among these, 218 genes were upregulated, while 177 genes were downregulated compared with the WT **(Fig. 4A and Table S3)**. To investigate the functional differences between WT and Δ*hsdMSR*, GO and KEGG databases were used to perform functional enrichment analyses based on the DEGs. When using adjusted p-value < 0.05 as a significant cut-off, the upregulated functional terms included alginate biosynthesis, fructose and mannose metabolism, and the oxidation– reduction process **(Fig. 4B)**. The downregulated pathways included the citrate cycle, oxidative phosphorylation, ribosome structure, and translation **(Fig. 4B)**. A total of 116 genes with bigger differences (|Log_2_FC| > 2) except for genes related to ribosomal protein, T3SS, and alginate synthesis. Among these genes, 31 were annotated as hypothetical proteins and 4 as transcription factors with unknown functions, and the remaining genes mostly encoded metabolism-related enzymes. These enzymes might have effects on growth defects in Δ*hsdMSR*.

**Figure 4.**
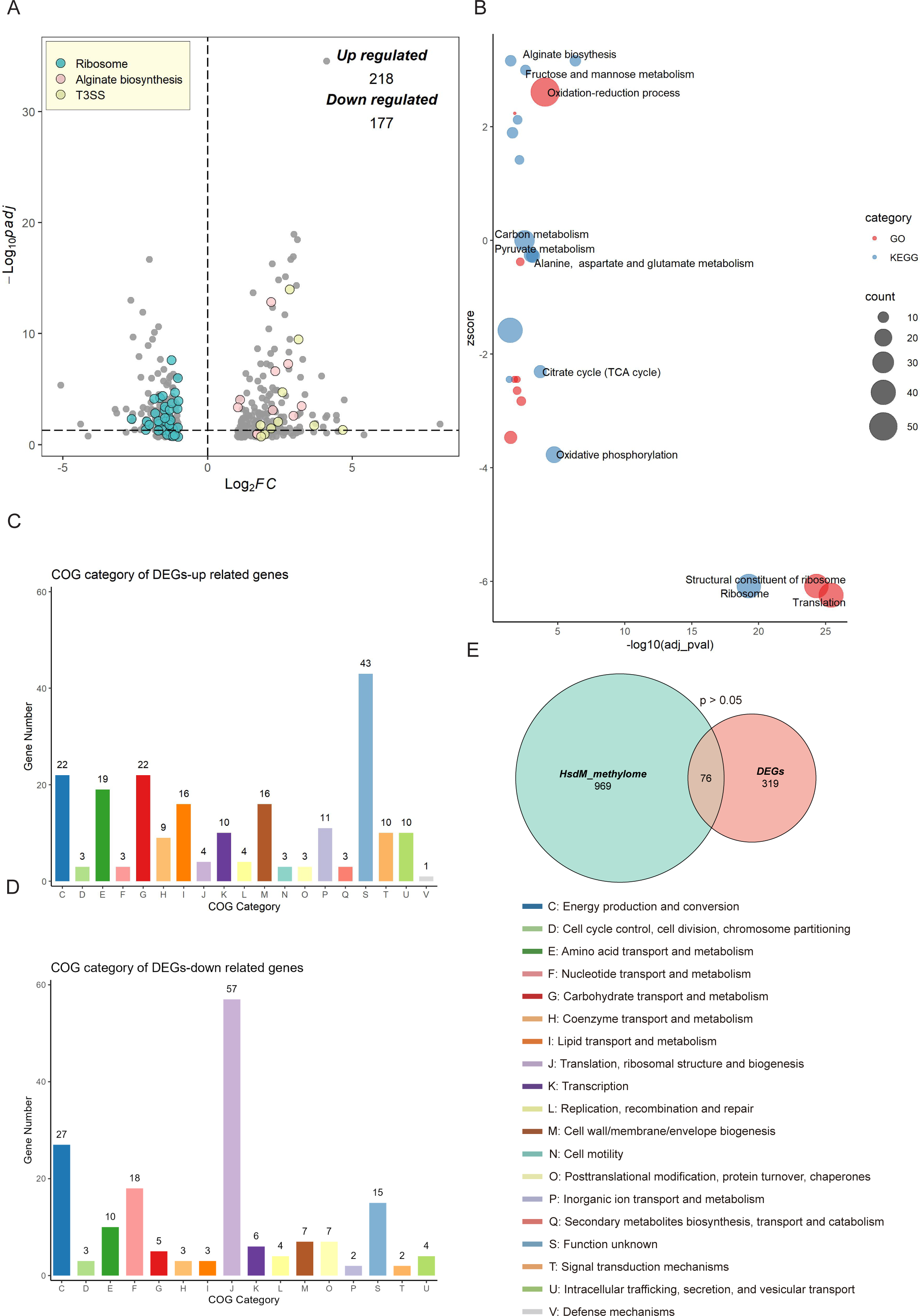
Transcriptional changes profiling of *hsdMSR* mutant in *Psph*. **(A)** The volcano plot reveals DEGs between *Psph* WT and Δ*hsdMSR*. The DEGs were (|log_2_ FC| > 1 and adjusted *P* value[<[0.05) in blue (Ribosomal protein), pink (Alginate biosynthesis), and yellow (T3SS). Each dot represents one gene. **(B)** Bubble plot shows enriched GO (Red) and KEGG (Blue) terms based on the DEGs between *Psph* WT and Δ*hsdMSR*. The x-axis shows the significance of functional annotation terms (−log10 adjusted *P*-value), and the y-axis indicates the Z-score of terms. The bubble size represents the gene number of terms. **(C)** COG classification and distribution of up-regulated DEGs. **(D)** COG classification and distribution of down-regulated DEGs. COG terms are highlighted in different colors. **(E)** Venn plot reveals the overlapped genes between DEGs and genes within the HsdMSR motif.

We also analysed the function alteration via COG category analysis of DEGs, which revealed that the upregulated genes were more likely to be involved in lanes E (amino acid transport and metabolism), G (carbohydrate transport and metabolism), and I (lipid transport and metabolism) **(Fig. 4C)**. Downregulated DEGs were significantly involved in lanes F (nucleotide transport and metabolism) and J (translation, ribosomal structure, and biogenesis) **(Fig. 4D)**. To investigate the correlation between DEGs and HsdM-recognising motif sites, we performed comparative analysis. Although the overlap was not statistically significant (p > 0.05), we identified 76 overlapping genes **(Fig. 4E)**. Eight DEGs harboured the HsdMSR methylation motif at the putative promoter region (within 100-bp upstream of the gene) **(Table S4)**.

### HsdMSR was required for T3SS and biofilm formation in *P. syringae*

To confirm the RNA-seq results, we performed quantitative real-time PCR (RT-qPCR) experiments on T3SS genes. Several T3SS effector genes were identified as upregulated, including *hopAE1*, *hrpF*, *hrpA2*, *hrpK1*, and *hrpW1*. RT-qPCR was applied under the same conditions as RNA-seq, and the transcriptional levels of all genes were confirmed as being significantly higher in the Δ*hsdMSR* than the WT **(Fig. 5A)**. These gene expression levels were restored in the complemented strain. To determine whether the loss of HsdMSR affected the *Psph* virulence phenotypes because of alterations to T3 effectors, we allowed the WT, Δ*hsdMSR*, and complemented strains to infiltrate the primary leaves of bean plants. At 6 days post-inoculation, Δ*hsdMSR* was observed to induce more severe symptoms than the WT and complemented strains **(Fig. 5B)**. Overall, HsdMSR inhibited the expression of T3 effectors under KB conditions, and the loss of this modification system enhanced the pathogenicity of the strain during plant infection.

**Figure 5.**
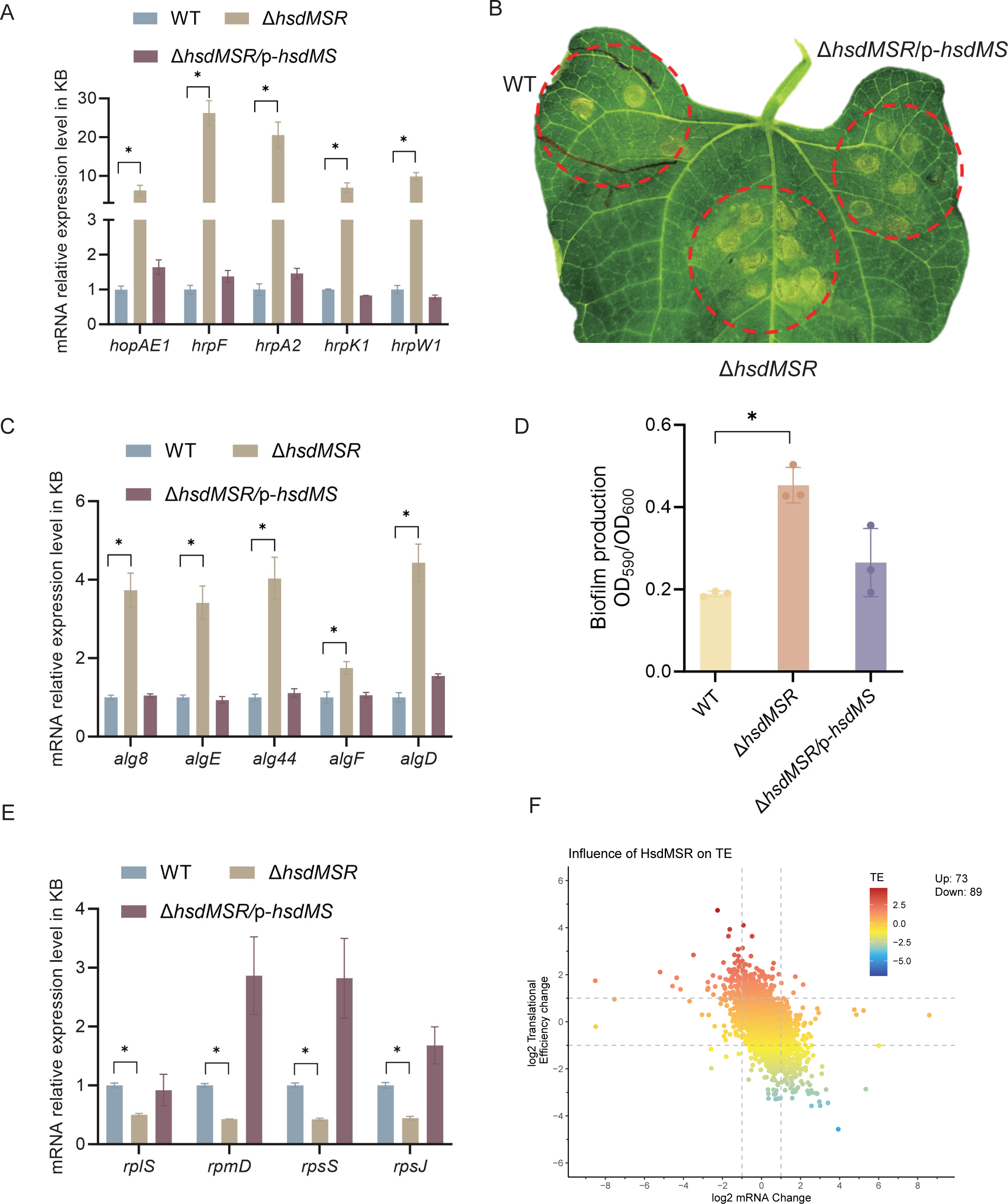
Influence of HsdMSR on virulence and metabolism in *Psph*. **(A)** HsdMSR negatively regulated T3SS-related genes. Data are shown as means ± SD (n = 3). **(B)** Disease symptoms caused by *Psph* WT, Δ*hsdMSR*, and the complemented strains, photographing 6 days after inoculation of 10^5^ CFU/mL bacteria. **(C)** HsdMSR negatively regulated alginate biosynthesis-related genes. Data are shown as means ± SD (n = 3). **(D)** The quantification of biofilm production in the *Psph* WT, Δ*hsdMSR*, and the complemented strains using a crystal violate staining assay. **(E)** HsdMSR positively regulated ribosomal protein-related genes. Data are shown as means ± SD (n = 3). **(F)** The scatterplot shows the TE and mRNA change between *Psph* WT and Δ*hsdMSR*. The x-axis presents the log2FC of the mRNA level, and the y-axis shows the log_2_FC of the TE. The greater TE is represented in red, whereas the lesser TE is displayed in blue.

Besides T3SS, alginate biosynthesis-related genes were also observed among the DEGs between WT and Δ*hsdMSR*. Alginate biosynthesis proteins have been demonstrated to be essential for Extracellular polymeric substances (EPS) production, which enhances biofilm formation during the bacterial infection process (47, 48). RT-qPCR experiments were performed on the relevant genes, including *alg8*, *algE*, *alg44*, *algF*, and *algD*. The expression levels of these genes in Δ*hsdMSR* were significantly increased compared with the WT and complemented strain **(Fig. 5C)**. To further investigate the influence of HsdMSR on biofilm formation, we performed a crystal violet staining assay to detect biofilm production by the three strains. The intensity of crystal violet staining of biofilms formed by the WT and complemented strains was significantly less strong than that of Δ*hsdMSR* cells **(Fig. 5D)**, supporting the role of HsdMSR in controlling biofilm formation. We therefore concluded that HsdMSR regulates the virulence of *P. syringae* by tuning its expression of T3SS and alginate biosynthesis genes.

### HsdMSR regulated ribosomal protein synthesis and translational efficiency

We noticed that HsdMSR was important for bacterial growth **(Fig. S4)**, while the expression of many ribosomal protein genes was reduced with the deletion of *hsdMSR* **(Fig. 5E)**. Regulation of ribosomal gene expression is essential to the integrity of the ribosome structure, which affects translation (49). Therefore, we hypothesised that the low expression level of ribosome proteins in Δ*hsdMSR* would result in delayed growth. We selected four genes encoding different ribosome subunits (*rplS*, *rpmD*, *rpsS,* and *rpsJ*) to verify the RNA-seq results through RT-qPCR. While *rplS* and *rpmD* encode 50S ribosomal proteins (L19 and L30), *rpsS* and *rpsJ* are responsible for 30S ribosomal proteins (S19 and S10). The RT-qPCR results demonstrated that the mRNA expression levels of these genes were significantly lower in Δ*hsdMSR* than in WT, while levels in the complemented strain were restored to the WT level **(Fig. 5E)**.

Given the important role of ribosomal proteins in translation, we performed ribosome profiling, also termed Ribo-seq, of the *Psph* WT and Δ*hsdMSR* strains to detect the influence of HsdMSR on translational efficiency (TE). When we compared the WT and Δ*hsdMSR*, the TE of 162 genes was significantly different (|log_2_FC| >1.5) (**Fig. 5F**). Of these genes, 73 genes had enhanced TE, while 89 had suppressed TE (**Fig. 5F**). Most of these genes were related to transmembrane transport and transcription regulation **(Fig. S6A-B)**. Remarkably, the TE of genes linked to the oxidation-reduction process tended to be suppressed **(Fig. S6B)**. These results showed that HsdMSR plays an important role in the regulation of metabolism by promoting ribosomal protein synthesis and modulating TE in *Psph*.

### HsdMSR regulation of *hrpF* was dependent on the full methylation of both strands

To investigate whether and how HsdMSR directly affects gene expression, we focused on those DEGs whose upstream regions harbour its motif. Interestingly, we noticed a methylation motif in the upstream region of *hrpF* (encoding the pathogenicity factor HrpF), which increased its expression level in the *Psph* WT strain **(Fig. 5A and 6A**). To further explore the effects of HsdMSR on the transcription of *hrpF*, we constructed a *lux*-reporter plasmid carrying the motif and extended it by 50 bp, which covered the upstream region of *hrpF*. After transferring the plasmid into the WT and Δ*hsdMSR* strains, we detected a significant difference in the expression level of *hrpF-lux* between these two strains. The *hrpF-lux* signal in Δ*hsdMSR* increased along with culture time and reached a peak at 24 hours when the methylation level was also elevated in the late growth stage **(Fig. 6B)**. This result of the *lux*-reporter assay confirmed the RNA-seq and RT-qPCR results showing that *hrpF* was remarkably upregulated in Δ*hsdMSR* compared with the WT strain.

**Figure 6.**
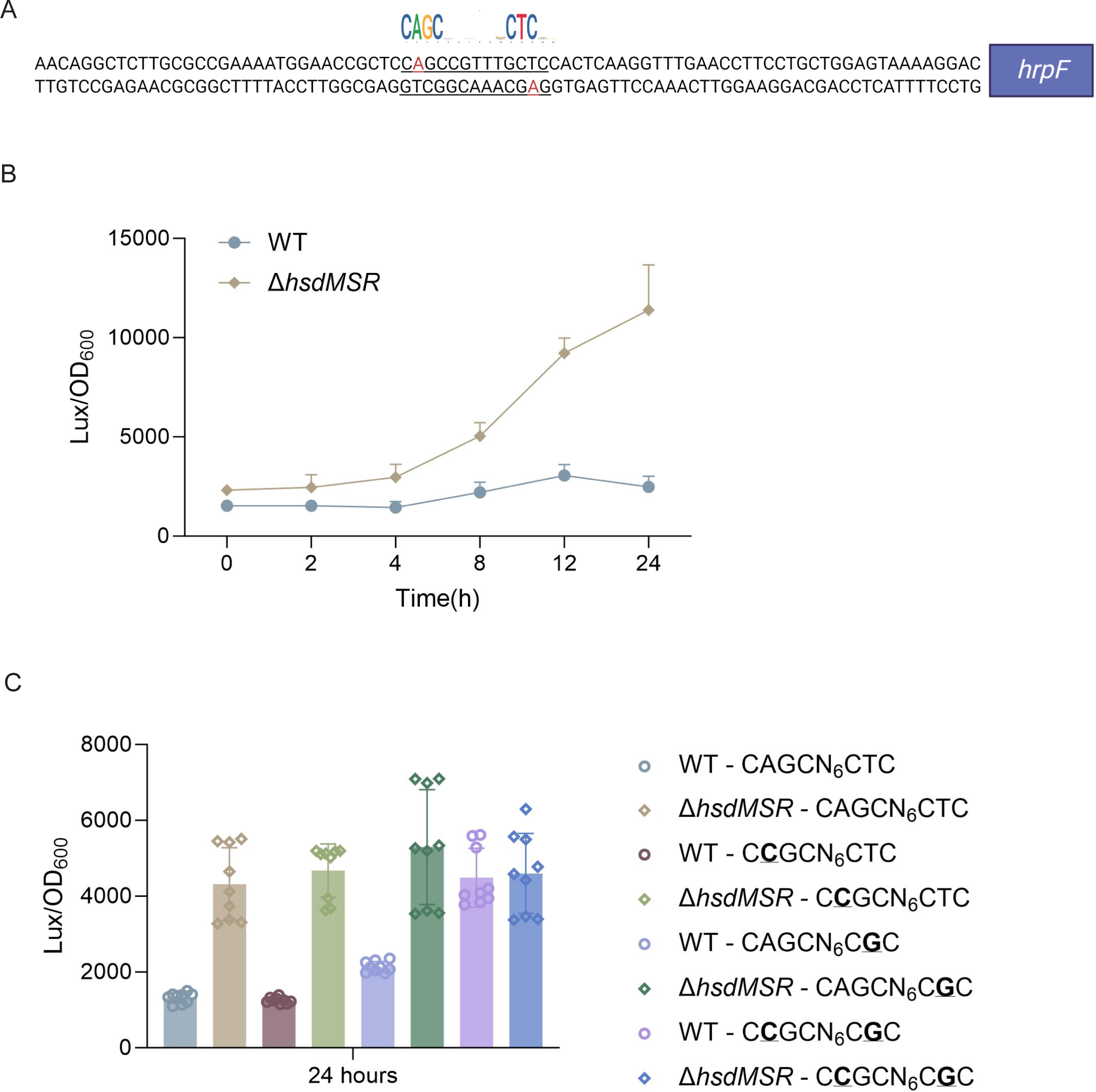
6mA methylation regulates gene transcription based on fully methylated. **(A)** Upstream region sequence of *hrpF* carrying the HsdMSR motif. Adenine methylation is highlighted in red. **(B)** Constantly lux activity detection of the *hrpF* between *Psph* WT and Δ*hsdMSR*. **(C)** Lux activity of *hrpF* between *Psph* WT and Δ*hsdMSR* with single- or double-point mutations. In the HsdMSR motif, “A” was replaced by “C” or “T” was replaced by “G”, highlighted in bold and underlined.

To detect the influence of the 6mA site within the HsdMSR motif on the expression of *hrpF*, we induced a point mutation on the single strand in the reporter plasmid to change the 6mA to C (C**<u>C</u>**GCN_6_CTC/ CCGCN_6_C**<u>G</u>**C). This resulted in a low *hrpF* expression level similar to that of the WT sequence carrying the A base **(Fig. 6C)**, indicating that a hemi-methylated pattern is insufficient for transcriptional alteration. To verify our hypothesis, we constructed a reporter with two A mutations (C**<u>C</u>**GCN_6_C**<u>G</u>**C), transferred it to the WT and Δ*hsdMSR* strains, and detected signal changes. As expected, the reporter carrying the HsdMSR motif without two A bases resulted in a significantly higher signal than the hemi-mutated and non-mutated reporters in the WT **(Fig. 6C)**. Taken together, the results showed that HsdMSR regulation of *hrpF* transcription was based on both strands being fully methylated.

## Discussion

Recent advances in sequencing technology have expanded our understanding of DNA methylation in bacteria. Numerous DNA MTases have been extensively characterised in different pathogenic bacteria, and data on their distribution patterns, recognition motifs, and biological functions have been collated (31, 32, 50–53). However, most research has concentrated on animal pathogenic bacteria, with limited insights into phytopathogenic bacteria except for *Xanthomonas*(54–56). Consequently, in this study, we employed SMRT-seq to profile the methylome patterns and specific conserved sequence motifs of three *P. syringae* strains. Notably, we discovered that the virulence and metabolism of *Psph* are regulated by Type I R-M MTase HsdM. The nonfunctional nature of the subunit restriction endonuclease HsdR was acknowledged, although its potential functionality relies on the presence of MTase (30).

Comprehensive analysis of the methylome atlas revealed that 6mA is the most prevalent modification type in *Psph* and *Pst*, mirroring the trends observed in other bacteria. Conversely, *Pss* displays the lowest occurrences of 6mA throughout its genome **(Fig. 1)**. This discrepancy might be attributed to the absence of the Type I R-M MTase responsible for 6mA in *Pss*. Similarly, the presence of two Type I R-M MTases in *Pst* possibly contribute to it having a higher number of 6mA sites than *Psph*. Additionally, all three pathovars had a similar pattern, with 4mC being the least frequent modification type. Despite belonging to the same *P. syringae* species, the three strains tested displayed remarkably divergent methylation patterns, which is reminiscent of the phenomenon observed in *Xanthomonas* spp. (54).

We further identified two (both 6mA) and four (three 6mA, one 4mC) motifs in *Psph* and *Pst*, respectively, but none in *Pss* **(Fig. 3)**. In fact, more motifs were predicted by the MEME and PacBio motif finders, but we chose only those motifs identified by both algorithms for higher accuracy. We found that the 6mA motifs had extremely high methylation levels, ranging from 77% to 100%, in the stationary phase of *P. syringae*. We subsequently found that the 6mA methylation levels increased during *Psph* growth, similar to the observations in *P. aeruginosa* (30). However, DNA methylation is much more stable under different conditions and growth phases in *Helicobacter pylori* and *Salmonella typhimurium* (57, 58).

MTases have been reported to play important roles in bacterial growth; for example, they are involved in cell cycle processes such as chromosomal replication in *E. coli* and *Caulobacter crescentus* (59–61). DNA methylation can activate DnaA and signal the initiation of chromosome replication to affect the growth of gamma-proteobacteria (62, 63). Previous studies revealed that DNA methylation influences bacterial growth through genes related to diverse carbohydrate transport mechanisms (55). We report that the 6mA modification catalysed by HsdM in *Psph* positively affects its growth, which can be explained by the downregulation of ribosomal proteins and the alteration of metabolism-related gene TE. However, many MTases are thought to be uncorrelated to growth in bacterial species such as *P. aeruginosa* and *Klebsiella pneumoniae* (30, 31, 52).

Many MTases and DNA methylations are involved in bacterial virulence through their modulation of gene expression (30, 31, 52, 53, 55, 56, 64–66). In this study, RNA-seq analysis of the HsdMSR mutation strain revealed 395 DEGs. Remarkably, not all DEGs harboured the HsdMSR methylation motif in their promoter region. There were no significant correlations between the HsdM-recognised motif sites in the promoter region and DEGs, even though DNA methylation is generally known to affect gene expression by altering the interactions between DNA and proteins such as TFs, which compete with MTases at specific motif sites and thus influence downstream transcription (67). However, recent studies have revealed that DNA methylation within CDS can also alter gene expression in bacteria; for example, 5mC in CDS can enhance transcription while blocking the transcription of CpG islands in promoters in eukaryotic cells (68, 69). We hypothesise that 6mA in *Psph* can regulate gene expression directly and indirectly. We confirmed that the 6mA motif in the promoter region of *hrpF* can directly and negatively regulate gene transcription in a manner dependent on full methylation of the strands **(Fig. 6)**. In addition, DNA methylation can change DNA curvature to decrease the thermodynamic stability of the double helix (70). Therefore, those DEGs carrying modified sites, including alginate biosynthesis-related genes and T3 effectors, are believed to be caused by the alteration of nucleoid topology, as seen in *Salmonella* and *H. pylori* (53, 71). Furthermore, the 6mA sites, including those in virulence-related *hrc/hrp* and *alg* genes, are conserved in *Psph* and *Pst*, implying that similar phenotypes occur in the latter strain. Altogether, 6mA modifications in *Psph* were observed to act as important epigenetic regulators of gene expression.

Apart from the established roles of 6mA and HsdMSR in *P. syringae*, certain signals or factors may influence HsdMSR expression. For instance, we confirmed that the growth phase affects methylation levels in *P. syringae*. Previous studies have shown that increased temperatures can reduce methylation levels, as observed in PAO1 (30). These findings suggest that HsdMSR expression may be responsive to both intrinsic cellular states and extrinsic environmental conditions. To further explore potential upstream TFs regulating the expression of HsdMSR, we searched for TF-binding sites in the HsdMSR promoter using our own databases (16, 18, 72). As a result, we found three candidate TFs (PSPPH_0061, PSPPH_3268, and PSPPH_3504) that might directly bind and regulate HsdMSR expression. Future studies on these TFs and their interactions with the HsdMSR promoter would help clarify the regulatory network governing HsdMSR activity.

Moreover, RM systems are known for their intrinsic role as innate immune systems in anti-phage infection, and present in around 90% of bacterial genomes (73). RM systems protect bacteria self by recognizing and degrading foreign phage DNA via methylation-specific site and restriction endonucleases (REases) (74). In addition, other phage-resistance systems are similar to RM systems but carry extra genes. One is called the phage growth limitation (Pgl) system, which modifies and cleaves phage DNA. However, the Pgl only modifies the phage DNA in the first infection cycle, and cleaves phage DNA in the subsequent infections, which gives a warn to the neighboring cells (75, 76). To counteract RM and RM-like systems, phages have evolved strategies, including unusual modifications such as hydroxymethylation, glycosylation, and glucosylation. They can also encode their own MTases to protect their DNA or employ strategies to evade restriction systems and other anti-RM defenses (77–79).

As more studies apply SMRT-seq to investigate bacterial methylomes, it has become evident that the epigenetic regulation of gene expression is highly prevalent among bacteria. Understanding the mechanisms through which methylation functions can provide novel insights into how strain-specific epigenetic modifications shape the adaptive responses of bacteria to distinct environmental challenges. As the repertoire of MTases varies among *P. syringae* strains, this approach will help us to better understand the diversity of DNA methylation and epigenetic patterns among *P. syringae* species.

## Materials and methods

### Bacterial strains and culture conditions

The bacterial strains, plasmids, and primers used in this study can be found in **Table S5**. All three model strains and mutations were cultured at 28[ in KB medium with shaking at 220 rpm or LB agar plates. The *E. coli* strains were grown at 37[ in LB broth (Luria-Bertani) with shaking at 220 rpm or on LB agar plates. Antibiotics were used at the following concentrations: kanamycin at 50 μg/mL, spectinomycin at 50 μg/mL, and rifampin 25 μg/mL.

### DNA extraction and SMRT sequencing

Three WT model strains and HsdMSR knockout *Psph* strains were cultured to the stationary phase. The genomic DNA of the strains was extracted using a TIANamp Bacteria DNA kit (Tiangen Biotech), using the manufacturer’s standard protocols. The SMRT sequence was performed at Abace Biotechnology company using Pacific Biosciences sequel II and Ile sequencer (PacBio, Menlo Park, CA, USA). SMRT-seq reads were aligned to the genome reference of *Psph* 1448A ( GCF_000012205.1), *Pst* DC3000 (GCF_000007805.1), and *Pss* B728a (GCF_000012245.1), respectively. SMRTLink software v13.0 was used to perform DNA methylation analysis. A modification quality value (QV) score of 50 and 100 was used to call the modified bases A and C, respectively.

### Deletion mutant and complemented strains construction

The restriction enzymes digested the pK18mobsacB suicide plasmid (80) listed in **Table S5**. The upstream (∼1500-bp) and downstream (∼1000-bp) fragments of HsdMSR open reading frame (ORF) were amplified from the *Psph* genome and digested. Then, the digested upstream and downstream fragments were ligated with T4 DNA ligase (NEB). The ligated fragments were inserted into the digested plasmid using ClonExpress MultiS One Step Cloning Kit (Vazyme). The recombinant plasmids were transformed into the *Psph* WT strain in the KB plate with rifampin and kanamycin. The single colonies were picked to a sucrose plate and then cultured in both KB with kanamycin/rifampin and KB with rifampin alone. Loss of kanamycin resistance indicated a double crossover. Finally, the deletion strains were further confirmed by qRT-PCR to detect the mRNA level of HsdMSR. For the complemented plasmids, the ORF of HsdMS was amplified by PCR from the *Psph* genome and cloned into the pHM1 plasmid.

### Growth curve measure assay

Overnight cultures of *Psph* and Δ*hsdMSR* were diluted to an OD_600_ of 0.1 in fresh KB liquid medium for use as the inoculum. One hundred μL of the inoculum was aliquoted into a 96-well microtiter plate in triplicate and incubated at 28°C for growth. OD_600_ values were recorded per 2 h for 8 times for plotting of growth curves.

### Dot blotting assay

Dot blotting was performed as previously described (81). Briefly, DNA samples were denatured at 95°C for 10 minutes and cooled down on ice for 3 minutes. Samples were spotted on the nylon membrane and air dry for 5 minutes, followed by heat-crosslink at 80[ for 2 hours. Membranes were blocked in 5% nonfat dry milk in TBST for 1 hour at room temperature, incubated with N^6^-mA antibodies (1:1000) overnight at 4°C. After 5 washes with TBST, membranes were incubated with HRP-linked secondary anti-rabbit IgG antibody (1:5,000) for 1 hour at room temperature. Signals were detected with ChemiDoc Imaging Systems (Bio-Rad). After imaging, incubate the membrane with methylene blue staining buffer for 15 minutes with gentle shaking.

### RNA-seq analyses

*Psph* and Δ*hsdMSR* strains were first cultured to the stationary phase in KB medium at 28°C. Cultures were collected, and total RNA was extracted using the RNeasy mini kit (Qiagen) following the manufacturer’s protocol and genomic DNA contamination was removed by DNaseI treatment (NEB). Subsequently, rRNA was depleted by using the MICROBExpress kit (Ambion), and the remaining mRNA was used to generate the cDNA library according to the NEBNext® UltraTM II RNA Library Prep Kit protocol (Illumina), which was then sequenced using the Illumina HiSeq 2000 system, generating 150-bp paired-end reads. Two biological replications have been performed. For each RNA-seq sample, raw sequencing reads were quality-trimmed using trim_galore (version 0.6.7) and aligned to the *Psph* genome using hisat2 (version 7.5.0)(82). DEGs were identified using DESeq2(83), and Function enrichment analysis of DEGs was conducted using the R package clusterProfiler(84).

### Ribo-seq library construction and analysis

The construction of the Ribo-seq library followed the previous protocol(85). Briefly, bacteria were cultured to an OD_600_ of 0.4, at which point chloramphenicol was added to a final concentration of 100 µg/mL for 2 minutes. Cells were then pelleted and washed with pre-chilled lysis buffer [25 mM Tris-HCl, pH 8.0; 25 mM NH_4_Cl; 10 mM MgOAc; 0.8% Triton X-100; 100 U/mL RNase-free DNase I; 0.3 U/mL Superase-In; 1.55 mM chloramphenicol; and 17 mM GMPPNP]. The pellet was resuspended in lysis buffer, followed by three freeze-thaw cycles using liquid nitrogen. Sodium deoxycholate was then added to a final concentration of 0.3% before centrifugation. The resulting supernatant was adjusted to 25 A_260_ units and mixed with 2 mL of 500 mM CaCl_2_ and 12 µL MNase, making up a total volume of 200 µL. After the digestion, the reaction was quenched with 2.5 mL of 500 mM EGTA. Monosomes were isolated using Sephacryl S400 MicroSpin columns, and RNA was purified using the miRNeasy Mini Kit (Qiagen). rRNA was removed using the NEBNext rRNA Depletion Kit, and the final library was constructed with the NEBNext Small RNA Library Prep Kit. For each sample, ribosome footprint reads were mapped to the *Psph* 1448A reference genome, and the translational efficiency was calculated by dividing the normalized Ribo-seq counts by the normalized RNA counts. Two biological replicates were performed for all experiments.

### Real-time quantitative PCR (RT-qPCR) verification

For RT-qPCR, RNA was purified using the RNeasy minikit (Qiagen). The cDNA synthesis was performed using a FastKing RT Kit (Tiangen Biotech). The assay was performed by SuperReal Premix Plus (SYBR Green) Kit (Tiangen Biotech) according to the manufacturer’s instruction. Each sample was repeated thrice with 600 ng cDNA and 16S rRNA as the internal control. The fold change represents the relative expression level of mRNA, which can be estimated by the threshold cycle (Ct) values of 2-(^ΔΔCt)^.

### Plant infection assay

The bean (*Phaseolus vulgaris* cv. *Red Kidney*) plants were used for the pathogenicity assay. The plant was grown in a greenhouse, as described previously (86). Overnight bacterial cultures were diluted to OD600 = 0.2 × 10^−3^ and were hand-inoculated into the primary leaves of week-old bean plants for 6 days.

### Biofilm formation assay

Biofilm production was detected, as previously reported (87). In brief, overnight bacterial cultures of *Psph* WT and HsdMSR mutants were diluted into OD600 = 0.1 and transferred to a 10 ml borosilicate tube containing 1 ml KB medium. Then, the cultures grow statically at 28°C for 3-5 days. Then, 0.1% crystal violet was used to stain the biofilm adhered to the tube tightly for 30 min, and other components bound to the tube loosely were washed off with distilled deionized water. The remaining crystal violet was fully dissolved in 1 ml 95%∼100% ethanol with constant shaking. Then, it was transferred to a transparent 96-well plate to measure its optical density at 590 nm using a Biotech microplate reader. The experiment was repeated using three independent bacterial cultures.

### Data analysis and statistics

The graphs in this paper were plotted using ggplot2 in R 4.2.0 software and GraphPad Prism 10.0.3 (GraphPad Inc.) Differences between groups were analyzed using students’ two-tailed t-tests. The results of all statistical analyses are shown as mean ± standard deviation (SD). All experiments were repeated independently at least three times with similar results.

## Data availability

The Ribo-seq, RNA-seq data, and SMRT-seq data were uploaded to the National Center for Biotechnology Information SRA database as part of BioProject PRJNA1055550 and PRJNA1123379, respectively. Codes were available upon reasonable request.

## Acknowledgments

This study was supported by grants from Guangdong Major Project of Basic and Applied Basic Research (2020B0301030005), Shenzhen Science and Technology Fund (JCYJ20210324134000002), the National Natural Science Foundation of China (32172358), General Research Funds of Hong Kong (11103221, 11101722, 11102223), the Sichuan Province Science and Technology Planning Project (2021YFSY0005). The funders had no role in study design, data collection, interpretation, or the decision to submit the work for publication.

## Author contribution

Conceptualization, J.H., and X.D; Methodology, J.H.; Validation, J.H., F.C., B.L., Y.Y., and Y.S.; Investigation, J.H., F.C. and B.L.; Writing & Editing, J.H., and X.D; Supervision X.D

## Declare of interest

The authors declare no competing interests.

## Supplementary information

**Figure S1. Distribution patterns of methylation sites in three model strains. (A)** Bar plot shows the distribution of modified sites located in different regions of CDS, intergenic region, and non-coding RNA. **(B)** GC contents distribution of 6mA between three *P. syringae* strains. **(C)** GC contents distribution of 5mC between three *P. syringae* strains. **(D)** GC contents distribution of 4mC between three *P. syringae* strains.

**Figure S2. COG analysis of methylation sites in *P. syringae*. (A)** COG classification of three DNA methylations in *Psph*. **(B)** COG classification of three DNA methylations in *Pst*. **(C)** COG classification of three DNA methylations in *Pss*. **Figure S3. Comparative genomic analysis of Type I RM system among bacterial species.** Different species are highlighted in different colors. The color key indicates the %identity compared to the MTases in *Psph*.

**Figure S4. Identification of Type I DNA methyltransferase in *Psph*. (A)** Growth curve of *Psph* WT, Δ*hsdMSR*, and complemented strains at 28°C in KB medium. **(B)** HsdMSR can catalyze 6mA formation in *Psph*. **(C)** The circle map displays the distribution of 6mA, 5mC, and 5mC in the HsdMSR mutant. **(D)** The Venn plot reveals overlapped genes within three types of DNA methylation of HsdMSR mutant. (**D**) The Venn plot reveals overlapped 6mA sites among *Psph* WT, HsdMSR mutant, and the loci of the motif CAGCN(_6_)CTC. **(F)** SMRT-seq reveals HsdMSR catalyzed the 6mA motif CAGCN(_6_)CTC.

**Figure S5. Effects of growth phases on methylation in *P. syringae*. (A)** Dot blot results of three *P. syringae* strains in logarithmic and stationary phases. **(B)** Overexpression of HsdM showed a higher 6mA modification level than that in WT. **(C)** Overexpression of HsdM did not affect the growth of *Pst*.

**Figure S6. Number of genes with significant TE changes. (A)** The GO terms for the genes in which TE changed more than 1-FC. **(B)** The GO terms for the genes in which TE changed less than 1-FC.

**Table S1 Restriction-modification systems predicted in P. syringae.**

**Table S2 Modified genes conserved in three P. syringae strains.**

**Table S3 DEGs genes between *Psph* WT and Δ*hsdMSR*.**

**Table S4 DEGs carrying HsdMSR motif in their putative promoter regions. Table S5 Genes with changed TE between *Psph* WT and Δ*hsdMSR*.**

**Table S6 Strains, plasmids, and primers.**

## References

1. Bull CT, De Boer SH, Denny TP, Firrao G, Saux MF-L, Saddler GS, Scortichini M, Stead DE, Takikawa Y. 2010. Comprehensive list of names of plant pathogenic bacteria, 1980-2007. Journal of Plant Pathology:551–592.

2. Xin X-F, Kvitko B, He SY. 2018. *Pseudomonas syringae*: what it takes to be a pathogen. Nature Reviews Microbiology 16:316–328.

3. Arnold DL, Lovell HC, Jackson RW, Mansfield JW. 2011. *Pseudomonas syringae* pv. *phaseolicola*: from ‘has bean’ to supermodel. Molecular Plant Pathology 12:617–627.

4. Xin XF, He SY. 2013. *Pseudomonas syringae* pv. *tomato* DC3000: a model pathogen for probing disease susceptibility and hormone signaling in plants. Annu Rev Phytopathol 51:473–98.

5. Monier JM, Lindow SE. 2005. Aggregates of resident bacteria facilitate survival of immigrant bacteria on leaf surfaces. Microb Ecol 49:343–52.

6. Clarke CR, Cai R, Studholme DJ, Guttman DS, Vinatzer BA. 2010. *Pseudomonas syringae* strains naturally lacking the classical *P. syringae* hrp/hrc Locus are common leaf colonizers equipped with an atypical type III secretion system. Mol Plant Microbe Interact 23:198–210.

7. Rahme LG, Mindrinos MN, Panopoulos NJ. 1992. Plant and environmental sensory signals control the expression of hrp genes in *Pseudomonas syringae* pv. *phaseolicola*. J Bacteriol 174:3499–507.

8. Xiao F, Goodwin SM, Xiao Y, Sun Z, Baker D, Tang X, Jenks MA, Zhou JM. 2004. *Arabidopsis* CYP86A2 represses *Pseudomonas syringae* type III genes and is required for cuticle development. EMBO J 23:2903–13.

9. Xiao Y, Lu Y, Heu S, Hutcheson SW. 1992. Organization and environmental regulation of the *Pseudomonas syringae* pv. *syringae* 61 hrp cluster. J Bacteriol 174:1734–41.

10. Xiao Y, Heu S, Yi J, Lu Y, Hutcheson SW. 1994. Identification of a putative alternate sigma factor and characterization of a multicomponent regulatory cascade controlling the expression of *Pseudomonas syringae* pv. *syringae* Pss61 *hrp* and *hrmA* genes. J Bacteriol 176:1025–36.

11. Hutcheson SW, Bretz J, Sussan T, Jin S, Pak K. 2001. Enhancer-binding proteins HrpR and HrpS interact to regulate hrp-encoded type III protein secretion in *Pseudomonas syringae* strains. J Bacteriol 183:5589–98.

12. Hendrickson EL, Guevera P, Ausubel FM. 2000. The alternative sigma factor RpoN is required for hrp activity in *Pseudomonas syringae* pv. *maculicola* and acts at the level of *hrpL* transcription. Journal of bacteriology 182:3508–3516.

13. Haefele DM, Lindow SE. 1987. Flagellar motility confers epiphytic fitness advantages upon *Pseudomonas syringae*. Applied and Environmental Microbiology 53:2528–2533.

14. Dasgupta N, Wolfgang MC, Goodman AL, Arora SK, Jyot J, Lory S, Ramphal R. 2003. A four-tiered transcriptional regulatory circuit controls flagellar biogenesis in *Pseudomonas aeruginosa* pv. *tomato* DC3000. Mol Microbiol 50:809–24.

15. Deng X, Liang H, Chen K, He C, Lan L, Tang X. 2014. Molecular mechanisms of two-component system RhpRS regulating type III secretion system in *Pseudomonas syringae*. Nucleic Acids Res 42:11472–86.

16. Fan L, Wang T, Hua C, Sun W, Li X, Grunwald L, Liu J, Wu N, Shao X, Yin Y, Yan J, Deng X. 2020. A compendium of DNA-binding specificities of transcription factors in Pseudomonas syringae. Nat Commun 11:4947.

17. Xie Y, Ding Y, Shao X, Yao C, Li J, Liu J, Deng X. 2021. Pseudomonas syringae senses polyphenols via phosphorelay crosstalk to inhibit virulence. EMBO Rep 22:e52805.

18. Shao X, Tan M, Xie Y, Yao C, Wang T, Huang H, Zhang Y, Ding Y, Liu J, Han L, Hua C, Wang X, Deng X. 2021. Integrated regulatory network in *Pseudomonas syringae* reveals dynamics of virulence. Cell Rep 34:108920.

19. Huang J, Yao C, Sun Y, Ji Q, Deng X. 2022. Virulence-related regulatory network of *Pseudomonas syringae*. Comput Struct Biotechnol J 20:6259–6270.

20. Gold M, Hurwitz J, Anders M. 1963. The enzymatic methylation of RNA and DNA, II. On the species specificity of the methylation enzymes. Proceedings of the National Academy of Sciences 50:164–169.

21. Casadesus J, Low D. 2006. Epigenetic gene regulation in the bacterial world. Microbiol Mol Biol Rev 70:830–56.

22. Wion D, Casadesus J. 2006. N6-methyl-adenine: an epigenetic signal for DNA-protein interactions. Nat Rev Microbiol 4:183–92.

23. Bickle TA, Kruger DH. 1993. Biology of DNA restriction. Microbiol Rev 57:434–50.

24. Loenen WA, Dryden DT, Raleigh EA, Wilson GG. 2014. Type I restriction enzymes and their relatives. Nucleic Acids Res 42:20–44.

25. Blow MJ, Clark TA, Daum CG, Deutschbauer AM, Fomenkov A, Fries R, Froula J, Kang DD, Malmstrom RR, Morgan RD, Posfai J, Singh K, Visel A, Wetmore K, Zhao Z, Rubin EM, Korlach J, Pennacchio LA, Roberts RJ. 2016. The Epigenomic Landscape of Prokaryotes. PLoS Genet 12:e1005854.

26. Beaulaurier J, Schadt EE, Fang G. 2019. Deciphering bacterial epigenomes using modern sequencing technologies. Nat Rev Genet 20:157–172.

27. Ho J, Taiaroa G, Butler MI, Poulter RTM. 2019. The Genome Sequence of M228, a Chinese Isolate of *Pseudomonas syringae* pv. actinidiae, Illustrates Insertion Sequence Element Mobility. Microbiol Resour Announc 8.

28. Poulter RTM, Ho J, Handley T, Taiaroa G, Butler MI. 2018. Comparison between complete genomes of an isolate of *Pseudomonas syringae* pv. actinidiae from Japan and a New Zealand isolate of the pandemic lineage. Scientific Reports 8.

29. Poulter R, Taiaroa G, Sumpter N, Stockwell P, Butler M. 2017. Complete Genome Sequence of the Kiwifruit Pathogen *Pseudomonas syringae* pv. actinidiae Biovar 5, Originating from Japan. Genome Announc 5.

30. Doberenz S, Eckweiler D, Reichert O, Jensen V, Bunk B, Sproer C, Kordes A, Frangipani E, Luong K, Korlach J, Heeb S, Overmann J, Kaever V, Haussler S.2017. Identification of a *Pseudomonas aeruginosa* PAO1 DNA Methyltransferase, Its Targets, and Physiological Roles. mBio 8.

31. Han S, Liu J, Li M, Zhang Y, Duan X, Zhang Y, Chen H, Cai Z, Yang L, Liu Y. 2022. DNA Methyltransferase Regulates Nitric Oxide Homeostasis and Virulence in a Chronically Adapted *Pseudomonas aeruginosa* Strain. mSystems 7:e0043422.

32. Li Z, Zhou X, Liao D, Liu R, Zhao X, Wang J, Zhong Q, Zeng Z, Peng Y, Tan Y, Yang Z. 2023. Comparative genomics and DNA methylation analysis of *Pseudomonas aeruginosa* clinical isolate PA3 by single-molecule real-time sequencing reveals new targets for antimicrobials. Front Cell Infect Microbiol 13:1180194.

33. Roberts RJ, Vincze T, Posfai J, Macelis D. 2023. REBASE: a database for DNA restriction and modification: enzymes, genes and genomes. Nucleic Acids Res 51:D629–D630.

34. Bird A. 2002. DNA methylation patterns and epigenetic memory. Genes & development 16:6–21.

35. Gelfman S, Cohen N, Yearim A, Ast G. 2013. DNA-methylation effect on cotranscriptional splicing is dependent on GC architecture of the exon–intron structure. Genome research 23:789–799.

36. Militello KT, Simon RD, Qureshi M, Maines R, Van Horne ML, Hennick SM, Jayakar SK, Pounder S. 2012. Conservation of Dcm-mediated cytosine DNA methylation in *Escherichia coli*. FEMS microbiology letters 328:78–85.

37. Nur I, Szyf M, Razin A, Glaser G, Rottem S, Razin S. 1985. Procaryotic and eucaryotic traits of DNA methylation in spiroplasmas (mycoplasmas). Journal of bacteriology 164:19–24.

38. Jin M, Chen J, Zhao X, Hu G, Wang H, Liu Z, Chen W-H. 2022. An engineered λ phage enables enhanced and strain-specific killing of enterohemorrhagic *Escherichia coli*. Microbiology Spectrum 10:e01271–22.

39. Ashburner M, Ball CA, Blake JA, Botstein D, Butler H, Cherry JM, Davis AP, Dolinski K, Dwight SS, Eppig JT, Harris MA, Hill DP, Issel-Tarver L, Kasarskis A, Lewis S, Matese JC, Richardson JE, Ringwald M, Rubin GM, Sherlock G. 2000. Gene ontology: tool for the unification of biology. The Gene Ontology Consortium. Nat Genet 25:25–9.

40. Kanehisa M, Goto S. 2000. KEGG: kyoto encyclopedia of genes and genomes. Nucleic Acids Res 28:27–30.

41. Cantalapiedra CP, Hernandez-Plaza A, Letunic I, Bork P, Huerta-Cepas J. 2021. eggNOG-mapper v2: Functional Annotation, Orthology Assignments, and Domain Prediction at the Metagenomic Scale. Mol Biol Evol 38:5825–5829.

42. Huerta-Cepas J, Szklarczyk D, Heller D, Hernandez-Plaza A, Forslund SK, Cook H, Mende DR, Letunic I, Rattei T, Jensen LJ, von Mering C, Bork P. 2019. eggNOG 5.0: a hierarchical, functionally and phylogenetically annotated orthology resource based on 5090 organisms and 2502 viruses. Nucleic Acids Res 47:D309–D314.

43. Bailey TL, Johnson J, Grant CE, Noble WS. 2015. The MEME Suite. Nucleic Acids Res 43:W39–49.

44. Gonzalez D, Collier J. 2013. DNA methylation by CcrM activates the transcription of two genes required for the division of *Caulobacter crescentus*. Mol Microbiol 88:203–18.

45. Krebes J, Morgan RD, Bunk B, Sproer C, Luong K, Parusel R, Anton BP, Konig C, Josenhans C, Overmann J, Roberts RJ, Korlach J, Suerbaum S. 2014. The complex methylome of the human gastric pathogen *Helicobacter pylori*. Nucleic Acids Res 42:2415–32.

46. Sanchez-Romero MA, Casadesus J. 2020. The bacterial epigenome. Nat Rev Microbiol 18:7–20.

47. Boyd A, Chakrabarty A. 1995. *Pseudomonas aeruginosa* biofilms: role of the alginate exopolysaccharide. Journal of industrial microbiology and biotechnology 15:162–168.

48. Ichinose Y, Taguchi F, Mukaihara T. 2013. Pathogenicity and virulence factors of *Pseudomonas syringae*. Journal of General Plant Pathology 79:285–296.

49. Zhou X, Liao WJ, Liao JM, Liao P, Lu H. 2015. Ribosomal proteins: functions beyond the ribosome. J Mol Cell Biol 7:92–104.

50. Murray IA, Clark TA, Morgan RD, Boitano M, Anton BP, Luong K, Fomenkov A, Turner SW, Korlach J, Roberts RJ. 2012. The methylomes of six bacteria. Nucleic Acids Res 40:11450–62.

51. Lee WC, Anton BP, Wang S, Baybayan P, Singh S, Ashby M, Chua EG, Tay CY, Thirriot F, Loke MF, Goh KL, Marshall BJ, Roberts RJ, Vadivelu J. 2015. The complete methylome of *Helicobacter pylori* UM032. BMC Genomics 16:424.

52. Fang CT, Yi WC, Shun CT, Tsai SF. 2017. DNA adenine methylation modulates pathogenicity of *Klebsiella pneumoniae* genotype K1. J Microbiol Immunol Infect 50:471–477.

53. Kumar S, Karmakar BC, Nagarajan D, Mukhopadhyay AK, Morgan RD, Rao DN. 2018. N4-cytosine DNA methylation regulates transcription and pathogenesis in *Helicobacter pylori*. Nucleic Acids Res 46:3815.

54. Seong HJ, Park HJ, Hong E, Lee SC, Sul WJ, Han SW. 2016. Methylome Analysis of Two *Xanthomonas* spp. Using Single-Molecule Real-Time Sequencing. Plant Pathol J 32:500–507.

55. Park HJ, Jung B, Lee J, Han SW. 2019. Functional characterization of a putative DNA methyltransferase, EadM, in *Xanthomonas axonopodis* pv. *glycines* by proteomic and phenotypic analyses. Sci Rep 9:2446.

56. Park HJ, Seong HJ, Lee J, Heo L, Sul WJ, Han SW. 2021. Two DNA Methyltransferases for Site-Specific 6mA and 5mC DNA Modification in *Xanthomonas euvesicatoria*. Front Plant Sci 12:621466.

57. Krebes J, Morgan RD, Bunk B, Spröer C, Luong K, Parusel R, Anton BP, König C, Josenhans C, Overmann J. 2014. The complex methylome of the human gastric pathogen *Helicobacter pylori*. Nucleic acids research 42:2415–2432.

58. Bourgeois JS, Anderson CE, Wang L, Modliszewski JL, Chen W, Schott BH, Devos N, Ko DC. 2022. Integration of the *Salmonella Typhimurium* Methylome and Transcriptome Reveals That DNA Methylation and Transcriptional Regulation Are Largely Decoupled under Virulence-Related Conditions. mBio 13:e0346421.

59. Reyes-Lamothe R, Sherratt DJ. 2019. The bacterial cell cycle, chromosome inheritance and cell growth. Nat Rev Microbiol 17:467–478.

60. Gonzalez D, Kozdon JB, McAdams HH, Shapiro L, Collier J. 2014. The functions of DNA methylation by CcrM in *Caulobacter crescentus*: a global approach. Nucleic Acids Res 42:3720–35.

61. Kozdon JB, Melfi MD, Luong K, Clark TA, Boitano M, Wang S, Zhou B, Gonzalez D, Collier J, Turner SW, Korlach J, Shapiro L, McAdams HH. 2013. Global methylation state at base-pair resolution of the *Caulobacter* genome throughout the cell cycle. Proceedings of the National Academy of Sciences 110:E4658–E4667.

62. Seong HJ, Han SW, Sul WJ. 2021. Prokaryotic DNA methylation and its functional roles. J Microbiol 59:242–248.

63. Boye E, Stokke T, Kleckner N, Skarstad K. 1996. Coordinating DNA replication initiation with cell growth: differential roles for DnaA and SeqA proteins. Proc Natl Acad Sci U S A 93:12206–11.

64. Manzer HS, Brunetti T, Doran KS. Identification of a DNA-cytosine methyltransferase that impacts global transcription to promote group B streptococcal vaginal colonization. mBio 0:e02306–23.

65. Banas JA, Biswas S, Zhu M. 2011. Effects of DNA methylation on expression of virulence genes in *Streptococcus* mutans. Appl Environ Microbiol 77:7236–42.

66. Oliveira PH, Ribis JW, Garrett EM, Trzilova D, Kim A, Sekulovic O, Mead EA, Pak T, Zhu S, Deikus G. 2020. Epigenomic characterization of *Clostridioides difficile* finds a conserved DNA methyltransferase that mediates sporulation and pathogenesis. Nature microbiology 5:166–180.

67. Lim HN, van Oudenaarden A. 2007. A multistep epigenetic switch enables the stable inheritance of DNA methylation states. Nat Genet 39:269–75.

68. Kahramanoglou C, Prieto AI, Khedkar S, Haase B, Gupta A, Benes V, Fraser GM, Luscombe NM, Seshasayee AS. 2012. Genomics of DNA cytosine methylation in *Escherichia coli* reveals its role in stationary phase transcription. Nat Commun 3:886.

69. Rishi V, Bhattacharya P, Chatterjee R, Rozenberg J, Zhao J, Glass K, Fitzgerald P, Vinson C. 2010. CpG methylation of half-CRE sequences creates C/EBPα binding sites that activate some tissue-specific genes. Proceedings of the National Academy of Sciences 107:20311–20316.

70. Diekmann S. 1987. DNA methylation can enhance or induce DNA curvature. The EMBO journal 6:4213–4217.

71. Camacho EM, Serna A, Madrid C, Marques S, Fernandez R, de la Cruz F, Juarez A, Casadesus J. 2005. Regulation of finP transcription by DNA adenine methylation in the virulence plasmid of *Salmonella enterica*. J Bacteriol 187:5691–9.

72. Sun Y, Li J, Huang J, Li S, Li Y, Lu B, Deng X. 2024. Architecture of genome-wide transcriptional regulatory network reveals dynamic functions and evolutionary trajectories in *Pseudomonas syringae*. bioRxiv:2024.01. 18.576191.

73. Oliveira PH, Touchon M, Rocha EP. 2014. The interplay of restriction-modification systems with mobile genetic elements and their prokaryotic hosts. Nucleic acids research 42:10618–10631.

74. Loenen WA, Dryden DT, Raleigh EA, Wilson GG, Murray NE. 2014. Highlights of the DNA cutters: a short history of the restriction enzymes. Nucleic acids research 42:3–19.

75. Hampton HG, Watson BN, Fineran PC. 2020. The arms race between bacteria and their phage foes. Nature 577:327–336.

76. Hoskisson PA, Sumby P, Smith MC. 2015. The phage growth limitation system in *Streptomyces coelicolor* A (3) 2 is a toxin/antitoxin system, comprising enzymes with DNA methyltransferase, protein kinase and ATPase activity. Virology 477:100–109.

77. Vasu K, Nagaraja V. 2013. Diverse functions of restriction-modification systems in addition to cellular defense. Microbiol Mol Biol Rev 77:53–72.

78. Murphy J, Mahony J, Ainsworth S, Nauta A, van Sinderen D. 2013. Bacteriophage orphan DNA methyltransferases: insights from their bacterial origin, function, and occurrence. Applied and environmental microbiology 79:7547–7555.

79. Iida S, Streiff MB, Bickle TA, Arber W. 1987. Two DNA antirestriction systems of bacteriophage P1, *darA*, and *darB*: characterization of *darA*− phages. Virology 157:156–166.

80. Kvitko BH, Collmer A. 2011. Construction of *Pseudomonas syringae* pv. *tomato* DC3000 mutant and polymutant strains. Methods Mol Biol 712:109–28.

81. Xie Q, Wu TP, Gimple RC, Li Z, Prager BC, Wu Q, Yu Y, Wang P, Wang Y, Gorkin DU. 2018. N6-methyladenine DNA modification in glioblastoma. Cell 175:1228–1243. e20.

82. Kim D, Langmead B, Salzberg SL. 2015. HISAT: a fast spliced aligner with low memory requirements. Nat Methods 12:357–60.

83. Love MI, Huber W, Anders S. 2014. Moderated estimation of fold change and dispersion for RNA-seq data with DESeq2. Genome biology 15:1–21.

84. Yu G, Wang L-G, Han Y, He Q-Y. 2012. clusterProfiler: an R package for comparing biological themes among gene clusters. Omics: a journal of integrative biology 16:284–287.

85. Hua C, Huang J, Wang T, Sun Y, Liu J, Huang L, Deng X. 2022. Bacterial Transcription Factors Bind to Coding Regions and Regulate Internal Cryptic Promoters. mBio 13:e0164322.

86. Xiao Y, Lan L, Yin C, Deng X, Baker D, Zhou JM, Tang X. 2007. Two-component sensor RhpS promotes induction of *Pseudomonas syringae* type III secretion system by repressing negative regulator RhpR. Mol Plant Microbe Interact 20:223–34.

87. Shao X, Xie Y, Zhang Y, Deng X. 2019. Biofilm Formation Assay in *Pseudomonas syringae*. Bio Protoc 9:e3237.

